# Exploring the Relationship between Geomagnetic Activity and Human Heart Rate Variability

**DOI:** 10.1101/684035

**Authors:** Matthew Mattoni, Sangtae Ahn, Carla Fröhlich, Flavio Fröhlich

## Abstract

Both geomagnetic and solar activity fluctuate over time and have been proposed to affect human physiology. One physiological measurement that has been previously investigated in this context, heart rate variability (HRV), has substantial health implications regarding the ability to adapt to stressors and has been shown to be altered in many cardiovascular and neurological disorders. Intriguingly, previous work found significant, strong correlations between HRV and geomagnetic/solar activity. In an attempt to replicate these findings, we simultaneously measured HRV from 20 healthy participants during a thirty-day period. In agreement with previous work, we found several significant correlations between HRV and geophysical time-series. However, after correction for autocorrelation, which is inherent in time-series, the only significant results were an increase in very low frequency during higher local geomagnetic activity and a geomagnetic anticipatory decrease in heart rate a day before higher global geomagnetic activity. Both correlations were very low. The loss of most significant effects after this correction suggests that previous findings may be a result of autocorrelation. A further note of caution is required since our and the previous studies in the field do not correct for multiple comparisons given the exploratory analysis strategy. We thus conclude that the effects of geomagnetic and solar activity are (if present) most likely of very small effect size and question the validity of the previous studies given the methodological concerns we have uncovered in our work.

## Introduction

Heart rate variability (HRV), an analysis of the change in the time intervals of consecutive heartbeats, is a well-established physiological measurement that serves as an indicator of disease and mortality risk ^1^. Previous studies have suggested that a higher degree of HRV is indicative of better health and lower risk for disease. For example, low HRV has been linked to myocardial infarction ^2^, neuropathy ^3^, depression ^4^, and schizophrenia ^5^. Diverse factors can modulate HRV, including genetic, neurological, respiratory, cardiovascular, lifestyle, and environmental factors ^6^. Analysis of HRV in the frequency domain is also used to estimate the activity of both the sympathetic and parasympathetic nervous system, though it has also been found to be dependent on heart rate and other confounding factors and thus possibly not necessarily a valid measure of autonomic activity ^7^.

Recent studies found that the change in the magnetic field of the earth caused by solar activity is significantly correlated with HRV ^8,9^. These studies were motivated by the relationship between cardiovascular health, specifically the occurrence of myocardial infarction, and both solar and geomagnetic activity ^10,11^. Yet, the relationship between HRV and geomagnetic activity remains unclear since several studies found conflicting results ^12,13^. Here, we attempted to replicate a previous study that showed strong and significant correlations between HRV and solar geomagnetic activity in a small pilot study ^8,9^. We found significant correlations between solar/geomagnetic activity and HRV components before correction for the autocorrelation inherent to time-series. However, we only found an increase in very low frequency HRV component and an anticipatory effect in heart rate with geomagnetic activity after correction for autocorrelation. Both effects were small; thus, previous studies have likely overestimated the effects due to the lack of stringent statistical analysis.

## Results

### Measurements of HRV and geomagnetic activity

We recruited 20 healthy participants and collected their 30-day (720 hours) longitudinal HRV data. Data from 19 participants were analyzed (1 dropout). We calculated heart rate (HR), standard deviation of NN intervals (SDNN), HRV triangular index (HRVTi), square root of the mean squared differences of successive RR intervals (RMSSD) as HRV time components. For HRV frequency components, very low-frequency power (VLF; 0.0033-0.04Hz), low-frequency power (LF; 0.04-0.15Hz), high-frequency power (HF; 0.15-0.4Hz), LF/HF ratio (LF/HF), and total power (VLF+LF+HF) were calculated. For measurements of geomagnetic activity, the K index and Ap index were obtained. In addition, F10.7 index was obtained as solar activity with 1-hour intervals.

### Correlations of HRV components

All HRV components were significantly correlated (Table 1, *p* < 0.001, both with and without the correction for autocorrelation). Measurements of overall variability were all positively correlated with each other and negatively correlated with heart rate. Each frequency-band percentage was inversely correlated to the others, an obvious relationship due to the shared denominator in their calculation. Total power was positively related to each measure of overall variability and HF_ratio_ (%), and negatively related to LF_ratio_ (%) and VLF_ratio_ (%), indicating that the HF band was responsible for the increase in the total power spectrum.

**Table 1.** Correlations between HRV components (\**p*<0.001, both with and without autocorrelation). HR: mean heart rate, VLF_ratio_: ratio of absolute VLF power to total power, LF_ratio_: ratio of absolute LF power to total power, HF_ratio_: ratio of absolute HF power to total power, LF/HF: ratio of absolute LF power to absolute HF power, total power: summation of absolute VLF, LF, and HF power. \**p*<0.001

|  | HR | SDNN | HRVTi | RMSSD | VLF<sub>ratio</sub> | LF<sub>ratio</sub> | HF<sub>ratio</sub> | LF/HF | Total Power |
| --- | --- | --- | --- | --- | --- | --- | --- | --- | --- |
| HR | 1 |  |  |  |  |  |  |  |  |
| SDNN | -.68* | 1 |  |  |  |  |  |  |  |
| HRVTi | -.67* | .91* | 1 |  |  |  |  |  |  |
| RMSSD | -.69* | .91* | .80* | 1 |  |  |  |  |  |
| VLF<sub>ratio</sub> | .37* | -.33* | -.36* | -.52* | 1 |  |  |  |  |
| LF<sub>ratio</sub> | .33* | -.34* | -.21* | -.36* | -.42* | 1 |  |  |  |
| HF<sub>ratio</sub> | -.64* | .61* | .53* | .82* | -.67* | -.39* | 1 |  |  |
| LF/HF | .58* | -.58* | -.50* | -.66* | .36* | .43* | -.72* | 1 |  |
| Total Power | -.59* | .94* | .80* | .89* | -.34* | -.30* | .59* | -.51* | 1 |

### Geomagnetic and solar activity

Ap index is a linear scale indicating global geomagnetic activity; values below 7 indicate a quiet period, values from 7 to 48 indicate an active or unsettled period, values from 48 to 80 indicate a minor storm, and values from 80 to 130 indicate a major storm. There were three notable geomagnetic events during the data collection period (Figure 1). The period 10/24-10/27 and 11/20-11/22 had small peaks in disturbance levels, peaking at 39 and 48 respectively, indicating periods of unsettled to a borderline minor storm of geomagnetic activity. A much larger event is noticeable in the 11/06-11/09 period, with a peak of 94 on 11/08, indicating that a major geomagnetic storm occurred. We also used the K index (Boulder magnetometer), which represents a semi-logarithmic 0-9 scale indicating local geomagnetic activity with 9 being the most activity. Kp index is an average of all global K indices, proving a semi-logarithmic measure of global activity. Values of each index varied from 0-6 during the time period of data collection, with the maximum of 6 occuring on 11/08, the same day as the Ap index maximum value (Figure 2). Unsurprisingly, the Ap index was strongly correlated to both the K index (*r*=.77, *p*<.001) and the Kp index (*r*=.86, *p*<.001); the K and Kp indices were also strongly correlated to each other (*r*=.85, *p*<.001). In addition, solar F10.7 index is presented, which exhibited the lowest value around 11/8 (Figure 1). F10.7 index stands for solar radio flux at wavelength of 10.7 cm and is a direct and reliable measure of solar activity ^14^. F10.7 is measured in solar flux units (sfu) and is often used as a proxy for other measures of solar activity.

**Figure 1:**
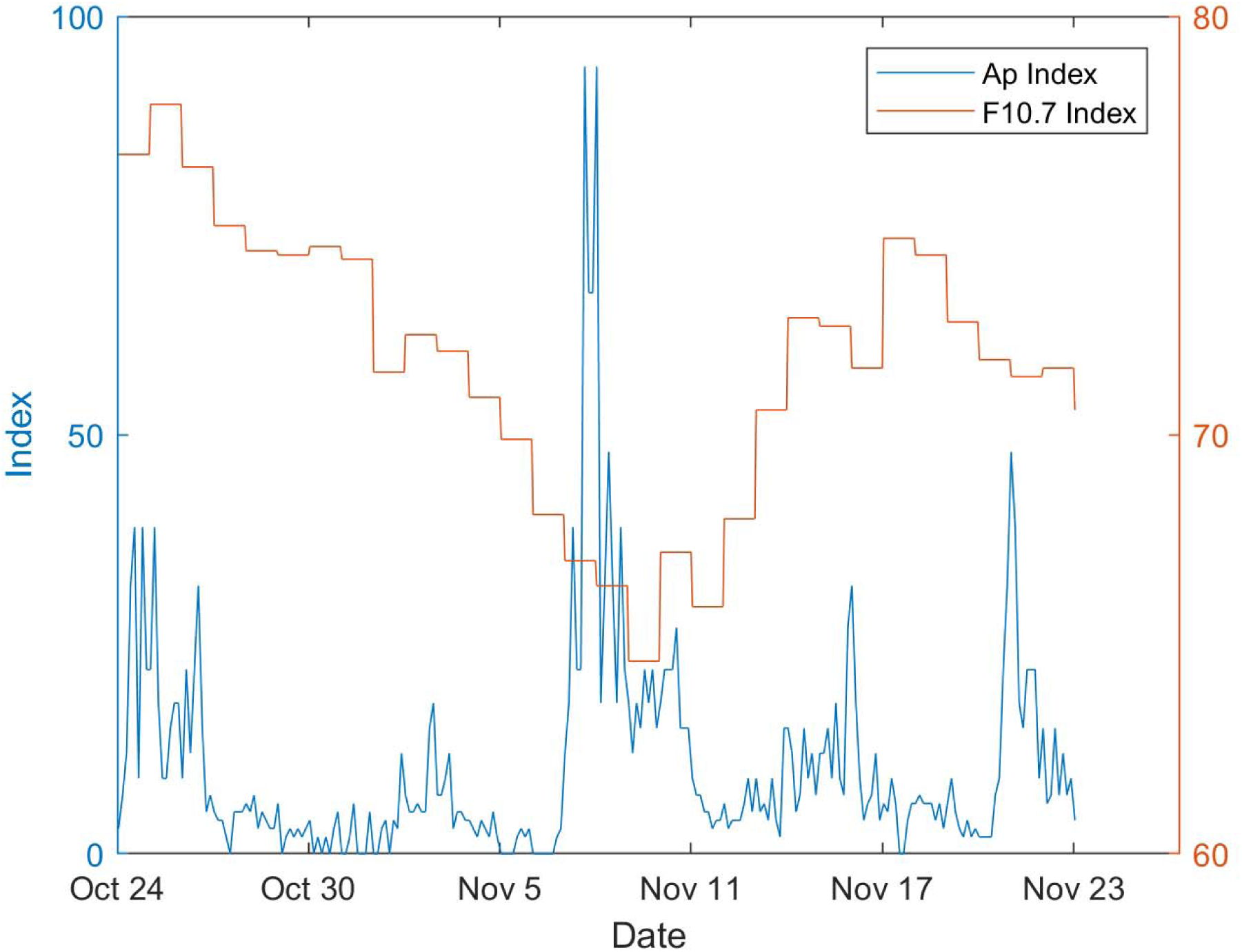
Geomagnetic Ap index and solar F10.7 index for the study period. See Methods Section for data sources.

**Figure 2:**
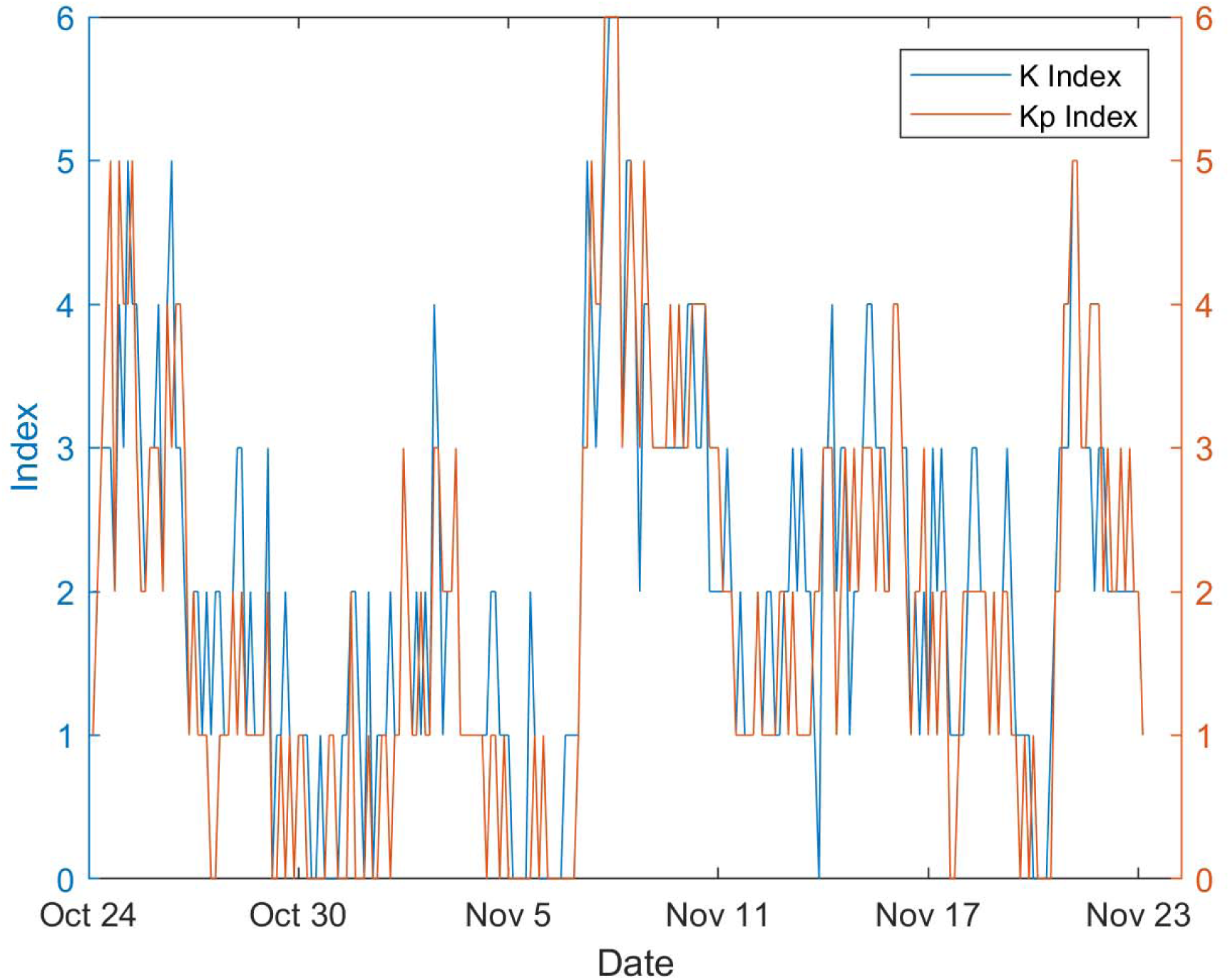
Geomagnetic K (Local, Boulder, CO) and Kp (Global) indices for the study period. See Methods Section for data sources.

### Relationship of HRV components with geomagnetic activity

Uncorrected and corrected for autocorrelation correlation coefficients between HRV components and geomagnetic activity are presented (Tables 2-4). We found significant correlations of Ap index with HRVTi, VLF_ratio_, LF_ratio_, and HF_ratio_ before correction but no significant correlations between Ap index and HRV components after correction (all *p*>0.05, Table 2). The K-index is significantly correlated with VLF_ratio_ and LF_ratio_ before correction, and the VLF_ratio_ correlation remained significant after correction, such that VLF_ratio_ increased as the local geomagnetic K-index increased (*r=*.06, *p*<.05, Table 3). We also found significant correlations of Kp-index with HR, RMSSD, VLF_ratio_, LF_ratio_, HF_ratio_, LF/HF, and total power before correction, but no significant correlations after correction (all *p*>0.05, Table 4).

**Table 2:** Correlation coefficients and *p*-values between HRV components and Ap index. Uncorrected and corrected p-values are presented. \**p*<0.05, \*\**p*<0.001

|  | Ap index |  |  |
| --- | --- | --- | --- |
| | $r$-value | $p$-value (uncorrected) | $p$-value (corrected) |
| HR | -.01 | .40 | .76 |
| SDNN | .01 | .34 | .65 |
| HRVTi | .03 | **.03\*** | .34 |
| RMSSD | -.01 | .42 | .74 |
| VLF<sub>ratio</sub> | .06 | **<.001\*\*** | .08 |
| LF<sub>ratio</sub> | -.03 | **.05\*** | .33 |
| HF<sub>ratio</sub> | -.03 | **.04\*** | .42 |
| LF/HF | .03 | .07 | .49 |
| Total power | -.01 | .40 | .68 |

**Table 3:** Correlation coefficients and *p*-values between HRV components and K index (Boulder, CO, magnetometer). Uncorrected and corrected p-values are presented. \**p*<0.05, \*\**p*<0.001

|  | K index |  |  |
| --- | --- | --- | --- |
| | r-value | $p$-value (uncorrected) | $p$-value (corrected) |
| HR | .02 | .25 | .63 |
| SDNN | -.01 | .93 | .77 |
| HRVTi | .02 | .30 | .54 |
| RMSSD | -.03 | .07 | .41 |
| VLF<sub>ratio</sub> | .06 | **<.001\*\*** | **.045\*** |
| LF<sub>ratio</sub> | -.04 | **.01\*** | .21 |
| HF<sub>ratio</sub> | -.03 | .08 | .44 |
| LF/HF | .01 | .34 | .78 |
| Total power | -.03 | .08 | .33 |

**Table 4:** Correlation coefficients and *p*-values between HRV components and Kp index. Uncorrected and corrected p-values are presented. \**p*<0.05, \*\**p*<0.001

|  | Kp index |  |  |
| --- | --- | --- | --- |
| | r-value | $p$-value (uncorrected) | $p$-value (corrected) |
| HR | .002 | **<.001\*\*** | .95 |
| SDNN | -.001 | .06 | .98 |
| HRVTi | .0001 | .30 | .98 |
| RMSSD | -.002 | **<.001\*\*** | .95 |
| VLF<sub>ratio</sub> | .002 | **<.001\*\*** | .95 |
| LF<sub>ratio</sub> | .001 | **.04\*** | .98 |
| HF<sub>ratio</sub> | -.003 | **<.001\*\*** | .95 |
| LF/HF | .002 | **<.001\*** | .96 |
| Total power | -.001 | **.01\*** | .97 |

A previous study ^8^ computed correlations after dividing the recording timeline into three distinct periods based on the occurrence of a geomagnetic storm, finding different relationships during the storm than before and after. To investigate this proposed phenomenon, we divided the Ap index values into three groups. The first group was the bottom 10^th^ percentile of Ap index values; all Ap index values here were 0, so this group was not further analyzed. The second group consisted of Ap index values between the 10^th^ and 90^th^ percentile (1-21), while the third group consisted of the top 90^th^ percentile (22-94). We calculated correlations between HRV components and AP index values for the two latter groups (Table 5). We found significant correlations between Ap index (top 90^th^ percentile) and HRVTi, Ap index (10^th^-90^th^ percentile) and HRVTi, VLF_ratio_, LF_ratio_, for uncorrected values. However, we found no significant correlations for corrected values.

**Table 5:** Correlation coefficients and *p*-values between HRV components and percentiles of Ap index. Uncorrected and corrected p-values are presented. \**p*<0.05

|  | Top 90<sup>th</sup> percentile |  |  | 10-90<sup>th</sup> percentile |  |  |
| --- | --- | --- | --- | --- | --- | --- |
| | $r$-value | $p$-value<br>(uncorrected) | $p$-value<br>(corrected) | $r$-value | $p$-value<br>(uncorrected) | $p$-value<br>(corrected) |
| HR | -.02 | .65 | .80 | .02 | .30 | .64 |
| SDNN | -.07 | .10 | .38 | .01 | .46 | .70 |
| HRVTi | -.08 | **.04\*** | .34 | .05 | **.01\*** | .18 |
| RMSSD | -.02 | .58 | .82 | -.01 | .47 | .72 |
| VLF<sub>ratio</sub> | .01 | .72 | .84 | .04 | **.02\*** | .21 |
| LF<sub>ratio</sub> | .00 | .93 | .84 | -.05 | **.01\*** | .20 |
| HF<sub>ratio</sub> | -.02 | .62 | .75 | .00 | .83 | .92 |
| LF/HF | .08 | .06 | .29 | .00 | .82 | .92 |
| Total power | -.06 | .13 | .56 | -.02 | .24 | .49 |

### Time-dependent relationship

To test for the potential presence of a time-lag between geomagnetic activity and change in HRV, we computed correlations between HRV components and corresponding Ap-index values 1 day after (anticipatory) and Ap-index values 1 day before (consequential). This analysis was motivated by previous studies ^8,9^ that discussed a potential “anticipatory effect”, which may relate to the fact that changes in solar activity take several days to modulate the geomagnetic field due to the time solar wind takes to reach the earth. In this analysis, we found a significant anticipatory relationship between heart rate and Ap index which survived the correction for autocorrelation of time-series; heart rate was negatively correlated with geomagnetic activity for corrected values (Table 6, *r*=-.09, *p*=.03). To further explore this significant correlation between hear rate and Ap index (anticipatory), we calculated correlations for each individual participant, again with and without correction for autocorrelation. Out of the 19 participants, 14 participants exhibited a negative correlation. In terms of significance testing, there were eight individuals with a significant correlation (all negative values) before correction for autocorrelation and three of these individuals exhibited a significant correlation after correction (Table 7).

**Table 6.** Time-dependent effect of Ap index with HRV components. Correlation coefficients and uncorrected and corrected *p*-values are presented. \**p*<0.05, **p<0.001

|  | Anticipatory |  |  | Consequential |  |  |
| --- | --- | --- | --- | --- | --- | --- |
| | $r$-value | $p$-value<br>(uncorrected) | $p$-value<br>(corrected) | $r$-value | $p$-value<br>(uncorrected) | $p$-value<br>(corrected) |
| HR | -.09 | **<.001\*\*** | **.03\*** | -0.02 | .70 | .89 |
| SDNN | .02 | .24 | .65 | -0.07 | .22 | .59 |
| HRVTi | .01 | .44 | .73 | -0.08 | **<.001\*\*** | .08 |
| RMSSD | .05 | **<.001\*\*** | .63 | -.02 | .14 | .81 |
| VLF<sub>ratio</sub> | -.03 | **.03\*** | .25 | 0.01 | .15 | .48 |
| LF<sub>ratio</sub> | -.03 | **.03\*** | .27 | .01 | .29 | .61 |
| HF<sub>ratio</sub> | .06 | **<.001\*\*** | .09 | -0.02 | **.02\*** | .35 |
| LF/HF | -.01 | .64 | .85 | 0.08 | .74 | .91 |
| Total power | -.003 | .84 | .92 | -0.06 | .62 | .80 |

**Table 7.** Individual anticipatory relationship of heart rate and Ap index. Demographics, correlation coefficients, and *p*-values are presented. \**p*<0.05, **p<0.001

| Participant # | Sex | Age | Anticipatory effect (Ap index and heart rate) |  |  |
| --- | --- | --- | --- | --- | --- |
|  |  |  | *r*-value | *p*-value<br>(uncorrected) | *p*-value<br>(corrected) |
| P01 | Male | 20 | .05 | .43 | .62 |
| P02 | Male | 20 | -.14 | **.04\*** | .23 |
| P03 | Female | 21 | -.25 | **<.01\*** | .08 |
| P04 | Female | 20 | -.11 | .13 | .37 |
| P05 | Male | 33 | -.01 | .97 | .98 |
| P06 | Female | 20 | -.39 | **<.001\*\*** | **.01\*** |
| P07 | Female | 19 | -.01 | .92 | .95 |
| P08 | Female | 21 | -.27 | **<.001\*\*** | **.03\*** |
| P09 | Female | 20 | -.14 | **.03\*** | .19 |
| P10 | Female | 18 | .07 | .28 | .56 |
| P11 | Female | 24 | -.17 | **<.01\*** | .14 |
| P12 | Female | 20 | .08 | .27 | .50 |
| P13 | Female | 48 | .03 | .70 | .80 |
| P14 | Female | 64 | -.16 | **.01\*** | .17 |
| P15 | Male | 37 | -.03 | .64 | .78 |
| P16 | Male | 23 | .04 | .50 | .69 |
| P17 | Excluded (please see section “Methods”) |  |  |  |  |
| P18 | Male | 25 | -.01 | .85 | .90 |
| P19 | Male | 31 | -.31 | **<.001\*\*** | **<.001\*\*** |
| P20 | Male | 31 | -.11 | .09 | .33 |

### Relationship of HRV with solar activity

For solar activity (F10.7 index), we found significant correlations between F10.7 index and SDNN, HRVTi, LF_ratio_, HF_power_, HF_ratio_. However, all significant effects were lost after correction for autocorrelation (Table 8). Of note, F10.7 index is a more direct measure of several solar processes and does not have a time-lag that Ap index may exhibit, which reflects changes in the geomagnetic field as a result of solar processes.

**Table 8.** Relationship of HRV with solar activity. Correlation coefficients and *p*-values are presented. \**p*<0.05, **p<0.001

|  | F10.7 index |  |  |
| --- | --- | --- | --- |
| | $r$-value | $p$-value (uncorrected) | $p$-value (corrected) |
| HR | .04 | **<.001\*\*** | .30 |
| SDNN | -.03 | **.01\*** | .48 |
| HRVTi | -.04 | **<.001\*\*** | .31 |
| RMSSD | -.01 | .39 | .85 |
| VLF<sub>ratio</sub> | -.05 | **<.001\*\*** | .11 |
| LF<sub>ratio</sub> | .04 | **<.001\*\*** | .25 |
| HF<sub>ratio</sub> | .02 | **.01\*** | .57 |
| LF/HF | -.01 | .37 | .84 |
| Total power | -.01 | .41 | .82 |

## Discussion

In this study, we investigated how geomagnetic/solar activity affects human HRV. We collected 720-hour HRV data from 19 participants and obtained geomagnetic and solar activity. Previous studies found significant, strong correlations between HRV and geomagnetic activity ^8,9^ In seeming agreement with this previous work, we also found significant correlations between the two type of variables before correction for the autocorrelation inherent to time-series. After correction for autocorrelation, however, we only found a significant correlation between very low-frequency power of HRV and K index and a significant anticipatory effect on heart rate with Ap index. Our results suggest that previous findings may be a consequence of autocorrelation instead of a true relationship between geomagnetism and HRV. We thus strongly recommend that correct statistical analyses should be performed when investigating this relationship.

Inspired by previous findings of time-dependent effects of geomagnetic activity on HRV ^8,12^, we also examined potential relationships with an offset of one day in both anticipatory or consequential manners. As a result, we found a significant relationship of an anticipatory effect such that heart rate was lower the day before higher geomagnetic activity, though the correlation was weak (r=-.09). Another study that examined time-dependent effects did not find this relationship to be significant ^12^. While this result possibly warrants further examination due to the various health abnormalities in which geomagnetism has been implicated ^12,15,16^, we note the possibility of a false positive considering the small effect size and the large amount of relationships tested to find this single result. For this effect to exist, the body would require a mechanism to predict geomagnetic storms. It is possible that the body detects abnormalities in geomagnetic fields before a sharp increase, or directly responds to changes in solar activity that reaches the earth before the factors that modulate the geomagnetic field. However, we would expect a significant relationship with solar activity to have existed if this was the case. Before further mechanistic speculation or examination, we recommend additional study replication, as this is the first study to find this specific effect.

As the intensity of geomagnetism varies with latitude, we also correlated HRV indices with the local geomagnetic K index from the Boulder magnetometer. After correction for autocorrelation, we found a relationship between the local K index and VLF_ratio_ such that the VLF band was stronger during stronger local geomagnetic activity. This finding may have clinical implications, as low VLF power has been more linked to all-cause mortality than the LF and HF bands ^17^, though we again note the weak correlation (r=.06) and the possibility of a false positive.

As a secondary focus, we report the effects of solar activity on HRV components. Similar to geomagnetic activity, solar activity has previously been correlated to SDNN, total power, LF, HF, VLF, and the LF/HF ratio ^8,9^. Again contrasting the results of previous studies, we found no significant relationships between HRV components and solar index F10.7. It is possible no significant effects were found due to the 20-participant sample size or the 1-month recording period. A study ^8^ reported significant relationships with very strong effect sizes, with some R^2^ values, such as the one between LF power and F10.7 index, reaching as high as 0.76. Results presented here vastly differ from this previous research, as we only found one significant relationship, which was small in magnitude. Noting the multitude of significant correlations we found before accounting for autocorrelation, the difference between this study and previous studies likely stems from the choice of the procedure to determine statistical significance. Time series correlated against each other may inherently be correlated as subsequent data points are dependent on each other, and the possible impact of this effect on studies examining environmental effects on HRV was pointed out in a Discover Magazine blog ^18^. Since all but one correlation lost significance after autocorrelation was removed, this study suggests that there is little to no relation between solar or geomagnetic activity and HRV, and contrasting findings are likely a result of the interdependence of data analyzed. It is unlikely that our test for autocorrelation erroneously removed significant relationships, as all correlations between HRV components (Table 1) survived the autocorrelation correction.

It is worth noting that there is an entire body of research on what is referred to as heliobiology. The full discussion of this literature is beyond the scope of this paper since many of the studies appear to exhibit serious flaws in methodology and reports ^19^. The potential mechanism of action remains speculative. In addition, there are other complicating factors in this research field, which include that the 11-year solar cycle makes studies hard to compare. For example, our study was performed near the nadir of the current solar cycle (near end of Cycle 24). Also, geographic latitude is likely to matter, and results may strongly depend on the latitude of the study site. Finally, it cannot be excluded that there are strongly differing levels of sensitivities across otherwise homogenous study populations. Our participant-by-participant analysis appears to support such individual differences in sensitivities to geomagnetic perturbations.

As any scientific study, our work has several limitations. First, larger sample sizes are desirable and warranted based on our results. Secondly, we tested numerous relationships and the few results that were significant at the p < .05 level were weak correlation values (r<.09), possibly indicating that the results are false positives. Thirdly, we have not collected any other psychological or biological variables which may explain the individual differences we found for the anticipatory relationship between heart rate and geomagnetic activity.

Overall, our study suggests that there is little to no effect of solar or geomagnetic activity on heart rate variability, with the only significant relationships being an anticipatory decrease in heart rate before increased global geomagnetic activity and an increase in very low-frequency power during periods of higher local geomagnetic activity. As in any research field, a single study does not provide final answers and more studies are warranted given the number of epidemiological studies that implicate solar/geomagnetic activity in human health.

## Methods

### Participants

We enrolled a total of 20 healthy participants over the age of 18 into this thirty-day longitudinal observational study. We recruited from the University of North Carolina at Chapel Hill area. Exclusion criteria included neurological or cardiovascular conditions, medication associated with these conditions, as well as pregnancy and daily meditation, which have been shown to affect HRV ^20,21^. This study was approved by the Biomedical Institutional Review Board of the University of North Carolina at Chapel Hill (IRB # 17-2397). All participants provided written informed consent before participating in the study. All methods were performed in accordance with the relevant guidelines and regulations.

### Materials

Each participant was provided with a commercially-available Firstbeat Bodyguard 2 (Firstbeat Technologies Oy, Jyväskylä, Finland) heart rate monitor and electrodes as well as the accompanying Firstbeat Uploader software (*https://www.firstbeat.com*). The monitor is worn on the torso with one electrode attached to the skin below the right collarbone, and the other electrode attached to the left ribcage; we instructed participants to move the electrode each day in a rotation of a few different spots in order to minimize skin irritation. The device automatically starts recording once both electrodes are attached, stores data internally, and has a battery life of approximately six days. Data were uploaded by participants through the Firstbeat software and REDCap (www.project-redcap.org), a secure online data collection portal for clinical research ^22^.

### Procedure

After recruitment, participants first uploaded a short sample recording to ensure that they were able to follow the study procedure. Once all these “practice samples” were obtained, participants were told the dates of the thirty-day data collection period, which was from October 24^th^, 2017 to November 22^nd^, 2017. Participants were instructed to begin wearing the monitor the night before the first day of the recording period. Participants were asked to wear the heart rate monitor nearly 24h/day and to only take it off for showering or other events that could cause water or other damage to the device. Participants uploaded data every 4 days; this pace allowed for participants to charge their device before the battery drained and for the research team to properly monitor data upload progress. If participants failed to consistently upload data, they were contacted by a member of the research team to provide data for the missing days. To upload their data, participants first loaded their data onto their computer through the Firstbeat uploader program and subsequently used a queue of surveys from REDCap for upload. In each survey, participants indicated the date(s) of the recording(s) and any time periods that they did not wear the monitor. They were also reminded to charge the monitor.

Data collection concluded after 30 days. At this time, we contacted participants to ensure that they had uploaded all collected data and to schedule a time to collect the device and provide compensation. To encourage participants to wear the monitor as often as possible, compensation included a flat $50 payment as well as a maximum bonus payment of $200 per participant. The amount of bonus a participant received was based on the amount of data provided, with compensation exponentially increasing with the amount of data provided.

### Environmental measurements

The K (Boulder) and Ap indices were obtained from the National Oceanic and Atmospheric Administration’s (NOAA) National Center for Environmental Information (*https://www.ncei.noaa.gov*) and are reported in 3-hour intervals. The K index was obtained from Kyoto University’s Data Analysis Center for Geomagnetism and Space Magnetism (http://wdc.kugi.kyoto-u.ac.jp/index.html). The F10.7 index was obtained from NASA’s Omniweb Data Explorer (*https://omniweb.gsfc.nasa.gov*) and is reported in 1-hour intervals.

### Data analysis

For 12 of the 20 participants analyzed, there were various clearly incorrect timestamps, which occurred when the devices reset to their factory restoration timestamp in 2015 for unknown reason. If possible, these times were corrected based on several identifiers, including relation to adjacent files, file upload timestamps, comparison between gaps in data and participant-identified times of not wearing the monitor, as well as other clues. Only files for which the correct time was determinable with high confidence were included (allowable margin of error: 15 minutes). Timestamps that could not be confidently corrected were not included in analysis. Eight participants had no errors in timestamps, nine participants had incorrect timestamps limited to the first five days or less, two participants had nearly half their data with incorrect timestamps, and one participant was not included in data analysis due to early withdrawal (reason: skin irritation). All time points for each participant were then concatenated into a single series and corrected to account for gaps in the time series created when participants did not wear the monitor. These times were finally adjusted to match UTC time in order to compare them to the environmental data.

RR interval data was first processed in Kubios HRV Premium, ver. 3.0.2 ^23^. The automatic artefact correction method in Kubios was used to correct for ectopic, too long, or too short beats by interpolating new RR values. Missed beats were corrected by adding new R-wave occurrence times and extra beats were corrected by removing extra R-wave detection and recalculating RR interval series. Further manual inspection of the RR series indicated that few artefacts were not removed and we thus removed RR values above 2.5s or below .2s.

Time and frequency domain analyses were completed in MATLAB R2016b (Mathworks, Natick, MA). Recordings were first split into 5-minute intervals as comparison of HRV between recordings of different lengths in not meaningful ^6^. Analysis was only completed for intervals containing data for at least 270 of the 300 seconds, and these results were then averaged into either 1-hour or 3-hour segments for comparison to environmental data. For time domain analysis, mean heart rate (HR), standard deviation of RR intervals (SDNN), HRV triangular index (HRVTi), and square root of the mean squared differences of successive RR intervals (RMSSD) indices were calculated. Standard deviation of average RR intervals (SDANN) results were not included, as this index is very similar to SDNN with 5-minute recordings. For frequency analysis, very low-frequency power (VLF; 0.0033-0.04Hz), low-frequency power (LF; 0.04-0.15Hz), high-frequency power (HF; 0.15-0.4Hz), LF/HF ratio (LF/HF), and total power (VLF+LF+HF) were obtained. Subsequently, a 24-hour moving average was used to remove circadian rhythms from HRV components.

We performed the Repeated measures correlation (RMC) analysis, a form of ANCOVA ^24^. RMC produces a common within-subject correlation effect between two variables. Overall, this method produces a single correlational value indicating how two variables are correlated. RMC was also used to correlate time-series of HRV components between participants. This allowed the HRV of participants to be compared to each other without a dependence on time, testing the synchronization of HRV over the duration of the measurement period.

### Correction for autocorrelation

When two time-series are being correlated with each other, the obtained results are possibly artificially inflated since individual samples of time-series are not independent due to inherent autocorrelation. To account for this, HRV data were stripped of their temporal structure by shuffling the data while keeping the original environmental data structure in place, and running RMC on this new dataset. Data with time points of 3-hours were shuffled in groups of 5 time points, and data with time points of 1-hour were shuffled in groups of 12 time points. This approach creates shuffled time-series that still exhibit most of the autocorrelation between neighboring samples but lack the temporal relationship with the other time-series it is correlated with. For each correlation, data were shuffled 1000 times, producing 1000 new RMC results. Empirical p values that have accounted for autocorrelation were then obtained with the equation:

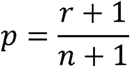

where,

*r* = amount of shuffled RMC values that are greater than the original RMC value

*n* = amount of shuffles (1000)

This method was used to calculate the p-value for every statistic presented. We report both raw and shuffle-corrected p-values since previous studies did not perform this correction for autocorrelation.

## Acknowledgments

This work was supported by the National Institute of Mental Health of the National Institutes of Health under Award Numbers R01MH111889 and R01MH101547, Psi Chi (The International Honor Society: Undergraduate Research Grant Fall), and Lindquist Undergraduate Research Fund. We gratefully acknowledge the help and support from the Carolina Center for Neurostimulation.

## Author Contributions

MM, CF, and FF designed the experiments; MM collected data; MM and SA analyzed the data; MM, SA, CF, and FF prepared the manuscript

